# BFVD - a large repository of predicted viral protein structures

**DOI:** 10.1101/2024.09.08.611582

**Authors:** Rachel Seongeun Kim, Eli Levy Karin, Martin Steinegger

## Abstract

The AlphaFold Protein Structure Database (AFDB) is the largest repository of accurately predicted structures with taxonomic labels. Despite providing predictions for over 214 million UniProt entries, the AFDB does not cover viral sequences, severely limiting their study. To bridge this gap, we created the Big Fantastic Virus Database (BFVD), a repository of 351,242 protein structures predicted by applying ColabFold to the viral sequence representatives of the UniRef30 clusters. BFVD holds a unique repertoire of protein structures as over 63% of its entries show no or low structural similarity to existing repositories. We demonstrate how BFVD substantially enhances the fraction of annotated bacteriophage proteins compared to sequence-based annotation using Bakta. In that, BFVD is on par with the AFDB, while holding nearly three orders of magnitude fewer structures. BFVD is an important virus-specific expansion to protein structure repositories, offering new opportunities to advance viral research. BFVD is freely available at https://bfvd.steineggerlab.workers.dev/

## Introduction

Viruses are infectious agents that invade host cells, exploiting their biological machinery for replication. They mutate rapidly, evading existing treatments and immunity, thus posing a persistent threat to public health (1). Their huge genetic diversity - often reflected in less than 30% amino acid sequence identity between newly discovered viruses and known ones, presents challenges for sequence-based annotation and classification. (2, 3). In contrast, due to their direct effect on function, protein structures tend to be more conserved, which can be used for studying viral mechanisms (4–6). Therefore, the availability of viral protein structures is critical for viral annotation through the detection of structural similarities.

Recent advancements in computational protein structure prediction (7–9) have made hundreds of millions of protein structures available through repositories like the AlphaFold Protein Structure Database (AFDB) (10, 11) and the ESM Atlas (8). These repositories have been transformative for studying the function of many proteins and protein families as a whole (12, 13). Using the AFDB has also contributed to the study of viruses, e.g., by improving the annotation of metagenomic bacteriophages (14) and by revealing viral proteins acquired from their metazoan host (15).

However, leveraging these vast resources for virus research remains limited. The AFDB excludes viral proteins (11), and the ESM Atlas lacks taxonomic information, making it difficult to identify viral proteins. Consequently, studying viral structures still relies on in-house prediction of protein structures (e.g., 16, 17), which is a time- and resource-consuming task. Recently, Nomburg et al. (18) presented ViralZone (19), a database of 67,715 predicted protein structures from 4,463 species of eukaryotic viruses. While offering an important milestone in the study of viruses, ViralZone remains focused on eukaryotic viruses and does not cover other viral clades.

Here, we focused on the viral fraction of UniProt (20) by examining its 30% sequence-identity clusters from UniRef30 (21, 22). We then predicted the protein structures of 351,242 cluster representatives of viral origin, consisting of over 3 million protein sequences. This effort resulted in BFVD, the largest repository of predicted viral structures to date. We show that BFVD contains highly diverse structures of various viral kingdoms, covering more viral variance than existing resources. We then demonstrate using BFVD for bacteriophage annotation, highlighting its utility tailored to viral research.

## Results

We first collected the representative protein sequences from UniRef30’s (21, 22) viral clusters, covering major viral clades (**Fig. 1a**). To limit the computational demand of structure prediction, we split 3,002 sequences longer than 1,500 residues (< 1% of all) into 6,730 sequence fragments. We then provided these fragments and the other sequences to ColabFold (23), resulting in 351,242 viral protein structures with a median predicted Local Distance Difference Test (pLDDT) of 70.17, indicating medium confidence (**Fig. 1b**). To assess the novelty of BFVD, we used Foldseek (24) to compare its structures to those of two major resources: AFDB50, a clustered version of the AFDB, consisting of 52 million cluster representatives (11) and the 100% sequence identity clustered Protein Data Bank (PDB100) with 279,193 entries (24, 25) **(Fig. 1c**). We found that ca. 15% of the structures were unique to BFVD, matching neither AFDB50, nor PDB100. An additional 9% matched only one of these databases. Furthermore, applying a cutoff on the quality of the match (TM score *≥*0.5), revealed that only 36% of the BFVD structures matched any of the two databases.

**Fig. 1.**
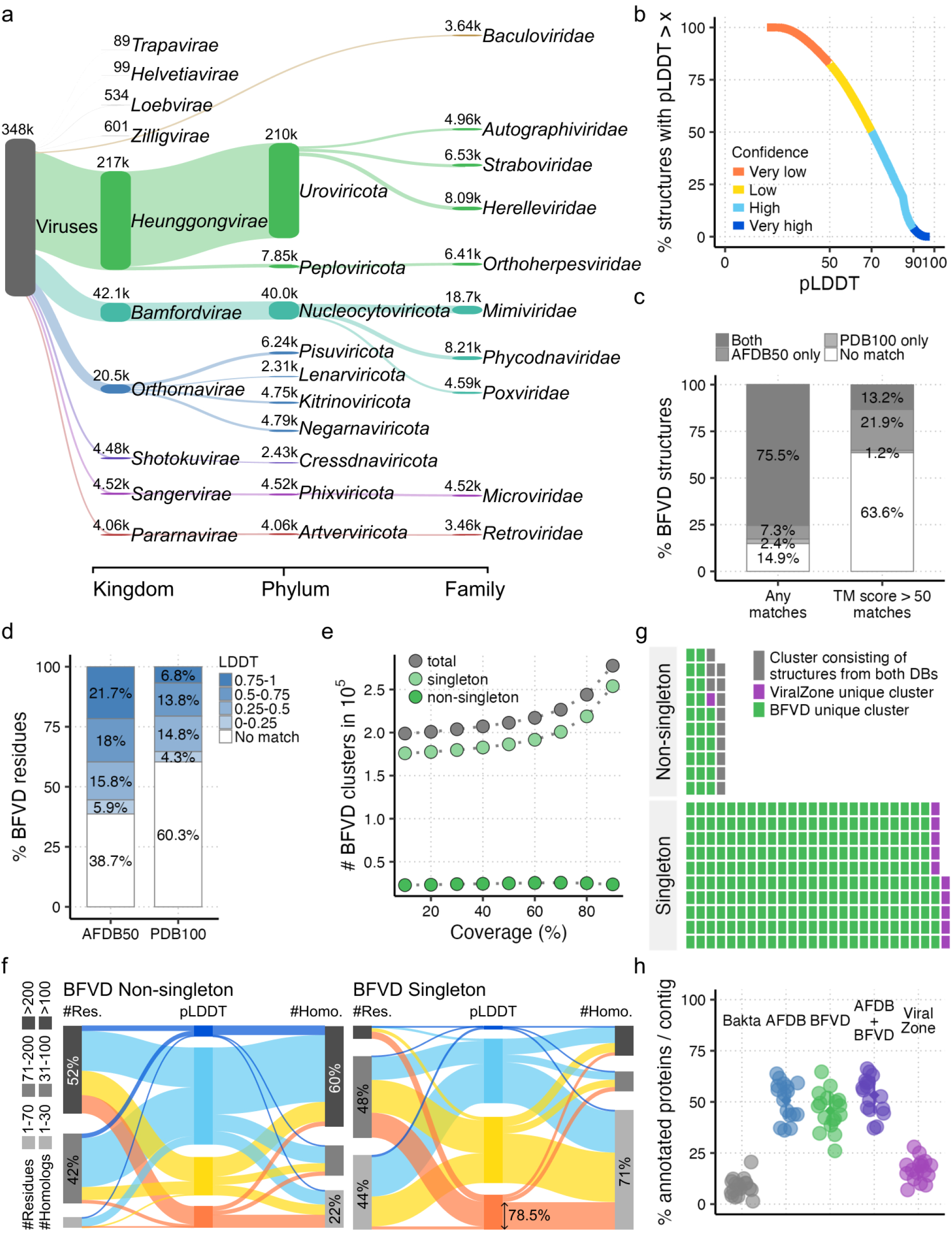
BFVD composition and comparison to other protein structure repositories. **a)** BFVD taxonomic composition. Shown at each rank are the 10 most abundant taxa. **b)** Cumulative distribution of pLDDT scores among BFVD’s 348k predicted structures. Over a half are highly confident. **c, d)** Foldseek comparison of BFVD to AFDB50 and PDB100 reveals its uniqueness. **c)** Ca. 15% of BFVD’s structures do not match AFDB50/PDB100 (left). Excluding low-similarity matches, ca. 64% of BFVD’s structures cannot be matched (right). **d)** Also on the residue-level, low similarity is observed against AFDB50 (left) and PDB100 (right). **e)** Structural redundancy reduction using Foldseek *cluster*. The number of clusters, especially singletons, increases with the value of the coverage parameter, though moderately until 70%. **f)** Examination of BFVD structures clustered at 70% coverage. Non-singleton proteins (left panel) are longer (left column) and have more homologs in their MSAs (right column) than singleton proteins (right panel). **g)** Joint clustering of BFVD and ViralZone. Each cell represents 800 clusters. BFVD’s structures are found in most clusters (96%), indicating its broad structural repertoire compared to ViralZone. **h)** Annotation of GAC6 contigs as in Say et al. Each point is a single contig, on top are the reference databases used for annotation. Sequence-reference (gray) annotates fewer proteins on each contig than structure-references (colors). BFVD annotates comparable fractions to the AFDB, while being much smaller.

We then evaluated residue-level similarity to these databases by retrieving the alignment LDDT values computed by Fold-seek for each match **(Fig. 1d**). We found that ca. 39% and 60% of the BFVD residues could not be aligned to AFDB50 and to PDB100, respectively. Additional fractions of ca. 6% and 4% could be matched to these databases only with a poor score (LDDT < 0.25). These results indicate that BFVD offers a unique opportunity to explore viral diversity that existing databases do not capture.

Next, we reduced structural redundancy in BFVD by using Foldseek *cluster* (13) to group together similar structures. We first studied the number of clusters obtained under different values of the Foldseek *cluster* coverage parameter (**Fig. 1e**). This parameter determines the minimal bidirectional coverage between a cluster representative and each cluster member, with lower values being more permissive. The total number of clusters increased from 198,868 to 277,631 following an increase in the coverage parameter. Furthermore, over 50% of the BFVD structures did not cluster and remained as singletons even at the lowest coverage threshold. In comparison, only 25% of the structures in AFDB50 (13 million out of 52.3 million) remained unclustered (13).

To investigate possible reasons so many BFVD protein structures failed to cluster, we focused on the clustering with 70% coverage cutoff, below which the number of singletons plateaued (**Fig. 1e**). We compared BFVD structures clustered as non-singletons (**Fig. 1f left**) and as singletons (**Fig. 1f right**) by three attributes: their lengths, the number of similar sequences (homologs) in their multiple sequence alignments (MSAs) used for structure prediction, and their pLDDT scores. We found that singleton proteins were shorter than non-singleton ones (median and average number of residues: 77 and 100, compared to 210 and 316). Focusing on the shortest proteins (*≤* 70 residues), we found that 99% of them were singletons. Unlike longer proteins, only 4% of the shortest proteins exhibited low confidence scores (pLDDT < 50). This is consistent with a previous report of high pLDDTs in sequences shorter than 100 residues (26).

Singleton protein structures also tended to have shallower MSAs, with an average of 254 homologs (median: 6), while non-singleton protein structures had an average of 1,274 (median: 183). As previously reported, prediction accuracy is negatively affected by an insufficient number of homologs, especially when there are fewer than 30 sequences (7, 27), leading to low confidence predictions. Indeed, among the low-confidence structure predictions (pLDDT *<* 50), the majority (78%) had fewer than 30 homologs. Put together, the high abundance of singletons in BFVD is likely driven by the short length and limited number of homologs of its proteins.

Recently, 67,715 protein structure predictions from eukaryotic viruses were made available in ViralZone (18, 19). To delineate the structural variation of BFVD and ViralZone, we applied Foldseek *cluster* to the joint set of their structures. This resulted in 31,965 non-singleton clusters, consisting of 211,645 structures, and in 204,323 singleton clusters (8,131 from ViralZone and 196,192 from BFVD) (**Fig. 1g**). Considering each of these 236,288 clusters as a putative structural class, we found that BFVD covered about 96% of all classes, by having a structure in nearly all (97%) non-singleton clusters and producing the most singletons. In contrast, ViralZone covered only 8% of all classes, by having a structure in 34% of all non-singleton clusters and producing substantially fewer singletons.

To demonstrate BFVD’s utility, we repeated and extended a part of a recent study by Say et al. (14) that annotated putative bacteriophages within metagenomically assembled contigs from wastewater. Say et al. developed a pipeline for enhanced annotations by integrating structural information from the AFDB with sequence data. Here, we applied the steps of their pipeline to one of the metagenomic samples from their study: the Granulated Activated Carbon sample 6 (GAC6). In addition to using the AFDB like they did, we included BFVD and ViralZone as reference databases for structural similarity search (**Fig. 1h**). Like Say et al., we found that the sequence-similarity based tool Bakta (28) could annotate on average 8% of the putative bacteriophage proteins on each contig, while Foldseek with the AFDB as reference annotated on average 51% of them. By using BFVD, we could annotate a comparable fraction of 46% of the putative bacteriophage proteins, despite the tremendous size difference between the AFDB and BFVD. However, when we searched the sample structures against the combined structure set of the AFDB and BFVD, we observed only a marginal increase in annotation performance. This suggests that the AFDB likely includes some BFVD bacteriophage structures indirectly, through prophages embedded in bacterial genomes covered by the AFDB. While ViralZone improved Bakta’s annotations, its contribution was limited compared to the AFDB and BFVD, likely due to its focus on eukaryotic viruses.

## Discussion

We presented BFVD, a database of 351k predicted proteins from the viral fraction of UniRef30. We showed that BFVD is unique and substantially different from the AFDB and the PDB as well as ViralZone. BFVD is far more comprehensive than ViralZone, as revealed by the analysis of their joint clustering. However, a notable finding was the high prevalence of singletons. As we showed, short proteins and shallow MSAs contribute to this phenomenon when clustering BFVD alone. Singletons should thus be treated with caution as they may not represent valid structural classes, but rather the result of poor structure prediction due to shallow MSAs or the presence of disordered regions. The utility of BFVD was demonstrated by achieving bacteriophage annotation performance comparable to the AFDB, effectively replacing the need for its 214 million entries with only 351k structures. This highlights the value of BFVD for virus-specific studies, offering a compact but comprehensive resource, tailored to their need. Moreover, since the entries in BFVD originate from UniProt, users can easily augment them with UniProt’s taxonomic and functional annotations to enhance the study of viral biology. BFVD’s structures can be used with current tools like Fold-seek as well as newly developed ones, like the multiple structure aligner FoldMason (29) to shed new light on viral function and evolution. Looking ahead, we aim to expand BFVD by predicting viral multimer structures, taking advantage of their compact genome size, and making them searchable using Foldseek Multimer (30). Ultimately, BFVD is an important resource that is expected to deepen our understanding of viral mechanisms from a structural perspective.

## Methods

### Preparing UniRef sequences for BFVD

The clustering at 30% pairwise sequence identity of UniProt (20) sequences release 2023_02 was downloaded from UniRef30 (21, 22) at https://gwdu111.gwdg.de/compbiol/uniclust/2023_02/. This dataset has 36,293,491 clusters, of which 347,514 had a viral sequence representative, as evident by their assigned taxnomic identifier (taxid), which is a descendant of taxid 10239. These clusters jointly contained 3,248,845 protein sequences and their 347,514 representatives were collected for the construction of BFVD. To limit the computational demand of the structure prediction step, we ensured that no sequence exceeded 1,500 residues in length. To that end, the 3,002 sequences longer than this threshold were split, resulting in 6,730 sequence fragments. BFVD was thus constructed from a total of 351,242 viral sequences.

### Taxonomic composition of BFVD

The taxid for each BFVD sequence was retrieved from UniProt and its full lineage - from NCBI (31). The Sankey plot based on this information (**Fig. 1a**) was generated with Pavian (32). For each taxonomic rank, only the ten most abundant taxa were included in the plot.

### Structure prediction

A multiple sequence alignment (MSA) was computed for each of the 351,242 sequences using MMseqs2 (version ede0be1) against the reference databases “uniref30_2022” and “colabfold_envdb_202108”. Based on these MSAs, structures were predicted using ColabFold v.1.5.2 (23) using the AlphaFold2 model and default parameters, except for ‘–num-models’ and ‘–stop-at-score’, which were set to 3 and 85, respectively. For each sequence, the best-ranking structural model according to the pLDDT score was kept.

### Exclusion of discontinuous structures

Out of the 351,242 structures predicted for BFVD, 3,259 (< 1%) contained the letter ‘X’ (unknown amino-acid) in their aminoacid sequence. As the presence of ‘X’ results in the prediction of discontinuous structures, we excluded these structures from further analysis, focusing on the other 347,983 structures in BFVD.

### Comparing BFVD to AFDB50 and PDB100

Foldseek (24) v.8.ef4e960 *easy-search* module was used to query the structures of BFVD against those of AFDB50 and of PDB100. The option ‘–greedy-best-hits’ was enabled to select only the best match for each BFVD query structure. For residue-wise assessment, the LDDT values computed by Foldseek for each alignment were extracted.

### Clustering

The redundancy reduction of BFVD was performed as described in “Results” using the Foldseek v.8.ef4e960 *easy-cluster* module. The same module was used with a coverage threshold of 0.7 for the joint clustering of 347,983 BFVD structures and 67,715 ViralZone structures. This coverage cutoff was selected based on Fig. 1d, as the point where the number of singletons plateaued.

### Bacteriophage annotation with BFVD

We followed the outline of the bacteriophage annotation pipeline described by Say et al. (14), with few modifications, as described in the following. Unlike Say et al., in the last step of the protocol, we added a search against BFVD and ViralZone.

#### Obtaining and assembling the GAC6 sample

The dataset for GAC6 was obtained from the European Nucleotide Archive accession PRJEB49151 (33) by selecting “t3_may7-2020” in the “sample title” field. The following steps were performed as described by Say et al., using the same parameters: base calling using Guppy (34) v.6.3.8, filtering using NanoFilt (35) v.2.8.0 and assembly using Flye (36) v.2.9.3-b1797. Unlike Say et al., we omitted the secondary assembly step and directly extracted circularized assemblies with a minimum coverage of 10.

#### Read mapping and polishing

Reads were mapped to each assembly using Minimap2 (37) v2.24, filtered by Gerenuq (38) v.0.2.3 and polished by Minipolish (39) v.0.1.3, using the same parameters as Say et al.

#### Bacteriophage detection

Following Say et al., the polished assemblies were annotated with Bakta (28) v.1.5.1 and then with INHERIT (40), retaining only assemblies annotated as bacteriophages.

#### Structure prediction

In all, 1,329 proteins were annotated as “bacteriophage” at the end of the last step. Like Say et al., we predicted the structures of these sequences using ColabFold v.1.5.5 with the same arguments. We retained for each sequence the best-ranking structural model according to the pLDDT score.

#### Foldseek search

The predicted structures were queried using Foldseek v8.ef4e960 *easy-search* module against the AFDB, like Say et al. In addition, they were queried against BFVD and ViralZone and the joint set of BFVD and AFDB.

## Acknowledgements

We thank Jaebeom Kim for assisting with formatting BFVD’s taxonomic information for the Sankey plot. We thank Milot Mirdita, Joe Grove and Uladzislau Litvin for their comments on the draft. M.S. acknowledges support by the National Research Foundation of Korea grants (2020M3-A9G7-103933, 2021-R1C1-C102065, 2021-M3A9-I4021220 and RS-2024-00396026), Samsung DS research fund, Creative-Pioneering Researchers Program and AI-Bio Research Grant through Seoul National University.

